# OTUB1 is a key regulator of RIG-I dependent immune signalling and is targeted for proteasomal degradation by influenza A NS1

**DOI:** 10.1101/634014

**Authors:** Akhee Sabiha Jahan, Elise Biquand, Raquel Muñoz-Moreno, Agathe Le Quang, Chris Ka-Pun Mok, Ho Him Wong, Qi Wen Teo, Sophie Doak, Alex W. H. Chin, Leo Lit Man Poon, Artejan te Velthuis, Adolfo García-Sastre, Caroline Demeret, Sumana Sanyal

## Abstract

Deubiquitylases (DUBs) regulate critical signaling pathways at the intersection of host innate immunity and viral pathogenesis. Although RIG-I activation is heavily dependent on ubiquitylation, DUBs that regulate this pathway have not been identified. Using a ubiquitin C-terminal electrophile, we profiled DUBs that function during influenza A virus (IAV) infection, and isolated OTUB1 as a key regulator of RIG-I dependent antiviral responses. OTUB1 was interferon-inducible, and interacted with RIG-I, viral PB2 and NS1. Upon infection, OTUB1 relocalised from the nucleus to mitochondrial membranes, and activated the RIG-I signaling complex via hydrolysis of K48 polyubiquitin chains and by forming a repressive complex with UBCH5c. Using a reconstituted system composed of in vitro translated [^35^S]IRF3, purified RIG-I, mitochondrial membranes and cytosol expressing OTUB1 variants, we recapitulated the mechanism of OTUB1-dependent RIG-I activation. A wide range of IAV NS1 proteins triggered proteasomal degradation of OTUB1, thereby antagonizing the RIG-I signaling cascade and antiviral responses.

**Highlights:**

1. OTUB1 is induced during influenza A virus infections in an IFN-I dependent manner
2. OTUB1 regulates the RIG-I complex by hydrolysing K48-linked polyubiquitin chains and by sequestering UBCH5c to prevent K48 polyubiquitylation
3. Optimal K63 versus K48 polyubiquitin chain concentrations determine RIG-I activation
4. Influenza NS1 targets OTUB1 for proteasomal degradation

---

Innate immunity against viral infections is triggered when germline-encoded pattern recognition receptors (PRRs) sense the presence of viral nucleic acids or other virus-specific components. The retinoic acid-inducible gene I (RIG-I)-like receptor (RLR) family constitutes one such group of cytosolic PRRs that is important in sensing viral RNA. Current evidence supports the activity of three members of the RLR family: RIG-I, melanoma differentiation-associated gene 5 (MDA-5), and laboratory of genetics and physiology 2 (LGP2). RIG-I and MDA-5 detect a wide range of RNA viruses, share several structural similarities, yet often display differences in a cell type and virus strain-specific manner (Jiang et al., 2012; Rehwinkel et al., 2010).

Much of viral sensing and immune response mechanisms derive from functional and structural studies of RIG-I (Yoneyama and Fujita, 2008). Upon binding to dsRNA, RIG-I transforms from an auto-repressed to an open conformation (Kowalinski et al., 2011); this facilitates its tetramerisation and K63-linked ubiquitylation by either TRIM25 or RIPLET (Cadena et al., 2019; Gack et al., 2007). RIG-I tetramers translocate to mitochondrial membranes (Liu et al., 2012) where they interact with mitochondrial antiviral signaling (MAVS) protein via their respective CARD domains. The resulting complex recruits downstream effector molecules (Gack, 2014; Pauli et al., 2014; Zeng et al., 2010), activates IRF3, IRF7 and NFκB to produce IFN-I, and creates an antiviral state.

Ubiquitylation, historically described for removal of misfolded proteins, has emerged as a critical regulator for fine-tuning signal transduction processes, and the RIG-I pathway is no exception. The mode of RIG-I ubiquitylation and its implications on signaling, has been the subject of intense research in the past few years. K63 ubiquitin chains, either in their free or conjugated forms, are critical for RIG-I signal transduction. Until recently K63 ubiquitylation was thought to be catalysed interchangeably by TRIM25 and RIPLET; however, recent studies have indicated that RIG-I activation is fundamentally dependent on RIPLET (Cadena et al., 2019).

All ubiquitin-dependent processes are accompanied by an additional layer of regulation by deubiquitylases (DUBs) (Komander et al., 2009). Many DUBs have been implicated in human diseases such as neurodegeneration, inflammation, infection, and cancer (Reyes-Turcu et al., 2009). Several DUBs, including A20 (Catrysse et al., 2014), CYLD (Trompouki et al., 2003), OTULIN, USP12 and Cezanne, have been described to function as regulators in NFκB and T-cell signalling (Duwel et al., 2009; Jahan et al., 2016), and are essential for resolving inflammation and restoring homeostasis. Despite the key role of ubiquitylation throughout the multi-step RIG-I signal cascade, the identities and function of DUBs that participate in the process are unkown.

In this study we applied activity-based profiling (Ovaa et al., 2004) to uncover DUBs that function during IAV infection, and identified OTUB1 as a critical regulator of RIG-I activation. We established that OTUB1 is key to optimal RIG-I signaling, and functions via a coordinated mechanism of hydrolysing K48 chains and forming an E2-repressive complex. CRISPR/Cas9-mediated knock-out of OTUB1 resulted in loss of oligomeric RIG-I, disassembly of the signaling complex and failure of NFκB and IRF activation in IAV-infected cells. We recapitulated this phenomenon in a reconstituted cell-free system to define the underlying mechanism of OTUB1 function in this pathway. Hydrolysis of K48 polyubiquitin chains and sequestration of UBCH5c by OTUB1 were prerequisites for K63 polyubiquitylation of RIG-I. Catalytically inactive OTUB1 displayed partial loss in IRF3 dimerisation due to the presence of redundant cytosolic DUBs. On the other hand, mutants that did not form the E2-repressive complex or inhibited OTUB1 redistribution to mitochondrial membranes abolished IRF3 activation. This phenomenon was antagonized by IAV NS1, which triggered proteasomal degradation of newly synthesized OTUB1. The underlying mechanisms that regulate access of high concentrations of K63 chains for RIG-I activation have not been studied. Our results therefore provide valuable insights into the mechanism of maintaining fidelity and optimal magnitude in the RIG-I signal transduction cascade, and highlight the complex interplay between immune responses and viral antagonism.

## Results

### Identification of host deubiquitylases activated upon influenza virus infection

Total cellular ubiquitylation profiles are often altered in IAV-infected cells, depending on severity of pathogenesis. To establish whether this occurs in a strain specific manner, we detected bulk ubiquitylation in IAV-infected A549 cells expressing wild-type ubiquitin. K48, K63 and M1-linked ubiquitylation profiles from mock and virus-infected cells were isolated by linkage specific tandem ubiquitin binding entities (TUBEs). While K48-linked ubiquitylation was consistently lower in infected cells, K63 ubiquitylated species were more pronounced in IAV-infected cells compared to mock. No significant changes were detectable in (M1) linear chains between the different IAV strains; however, they were elevated in infected cells as compared to mock. Bulk profiles therefore indicated distinct differences that exist in the dynamics of ubiquitylation upon infection by different strains of flu virus (Figure 1A).

**Figure 1.**
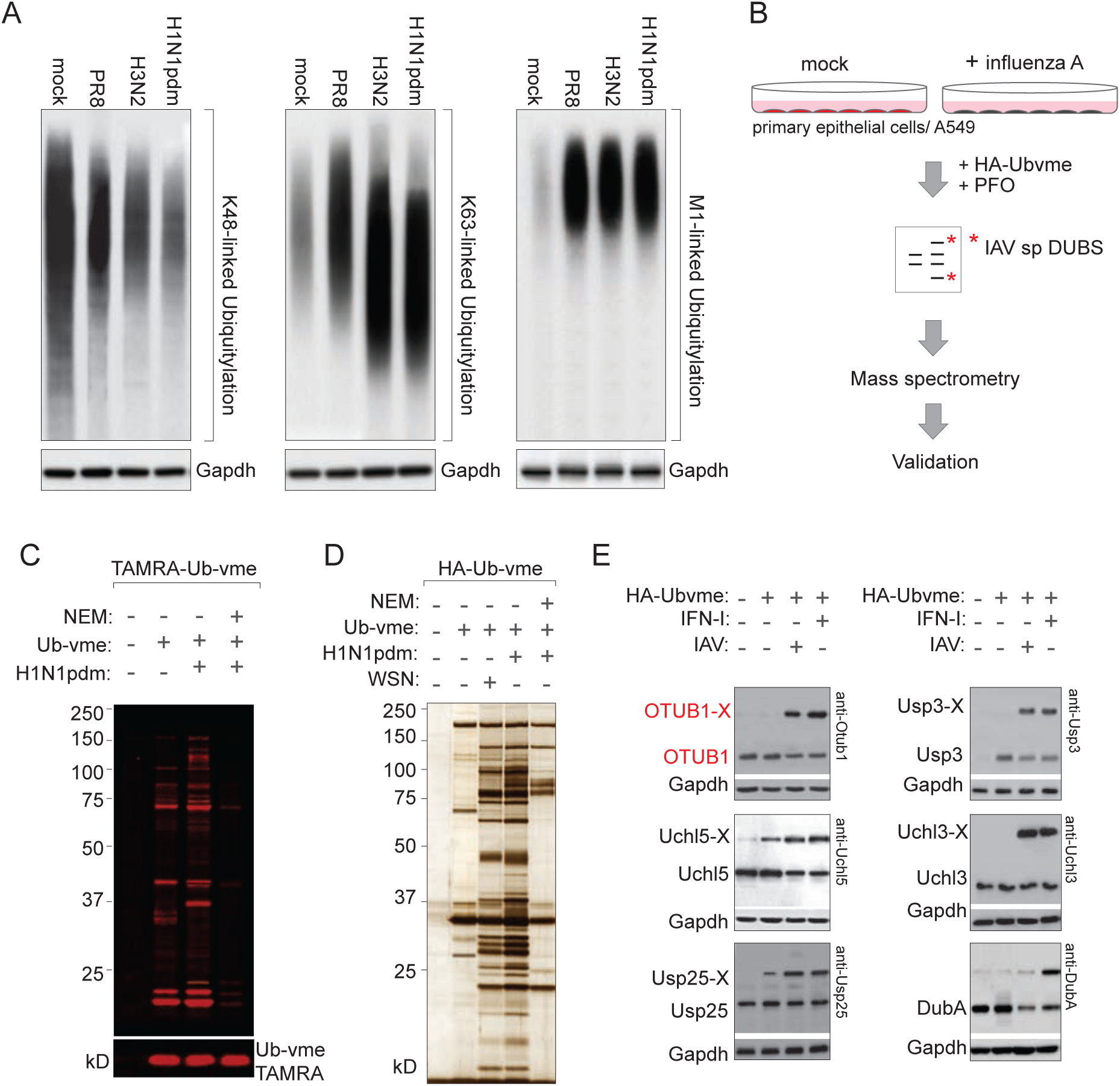
Identification of deubiquitylases activated upon influenza A virus infection. **(A)** Linkage-specific ubiquitylation profiles isolated from mock and IAV-infected samples. A549 cells expressing wild-type ubiquitin were infected with different strains of influenza A virus; A/PR/8/34, A/H1N1pdm09, A/Oklahoma/309/06 (H3N2). Ubiquitylated proteins were isolated on linkage specific tandem ubiquitin binding entities (TUBES), resolved by SDS-PAGE and visualised by linkage specific anti-ubiquitin antibodies; anti-K48, anti-K63, and anti-M1 linear chains. **(B)** Schematic for isolation of deubiquitylases activated upon influenza A virus infection. **(C)**, **(D)** Tamra and HA-tagged Ub-vme treated mock- and IAV-infected primary epithelial cells. Silver stained profile of large-scale immunoprecipitation of Ubvme-modified DUBs from mock and IAV-infected cells were subjected to mass spectrometry for protein identification. **(E)** Validation of identified proteins and their reactivity towards Ub-vme in IAV-infected or interferon-treated samples was verified by immunoblotting.

To isolate DUBs from IAV-infected cells, we followed an experimental set-up as illustrated in the schematic (Figure 1B) (Jahan et al., 2016; Zhang et al., 2018). We and others have previously employed similar approaches with ubiquitin C-terminal electrophiles to isolate cellular DUBs (Jahan et al., 2016; Kwasna et al., 2018).

Mock and H1N1 pandemic IAV-infected primary lung epithelial cells or A549 cells were permeabilised using perfringolysin O (Sanyal et al., 2013; 2012), and DUBs activated during infection were isolated with Ub-vme, either carrying a TAMRA or HA-tag. Ub-vme reactive material from control and infected cells were resolved by SDS-PAGE and visualised by fluorescent scanning (Figure 1C) or enriched on anti-HA beads first and detected by silver staining (Figure 1D). To preclude identification of non-specific or secondary interactors, cysteine-reactive N-ethyl maleimide treated samples prior to adding Ub-vme were included as a control. Potential candidates were identified by trypsin digestion, mass spectrometry and spectral counting of peptides on immunoprecipitated material (**Table S1**).

We validated the MS hits in lysates from mock and either virus-infected or IFN-I treated samples by immunoblotting, where we detected a shift in molecular weight in proteins that reacted with Ub-vme (Figure 1E). Among candidates identified were known regulators of interferon signaling, such as Usp25 (Lin et al., 2015), Usp15 (Pauli et al., 2014) and DUBA, which displayed reduced expression in H1N1-infected cells and high Ubvme reactivity in IFN-I treated samples, in agreement with a previous report (Kayagaki et al., 2007).

### OTUB1 is an interferon inducible host factor that is specifically degraded by influenza A virus

OTUB1 is a member of the Ovarian tumor domain containing superfamily of proteases. It has a catalytic cysteine (C91), several phosphorylation sites (phosphorylation on S16 can determine cytosolic versus nuclear localisation of OTUB1 (Herhaus et al., 2014)), and hydrolyses K48 ubiquitin chains (Edelmann et al., 2009; Juang et al., 2012) (Figure 2A). It is also able to inhibit ubiquitylation via its non-canonical activity of sequestering E2 enzymes (Juang et al., 2012). While expressed is most tissues, it displays particularly high mRNA levels in lung epithelial cells.

**Figure 2.**
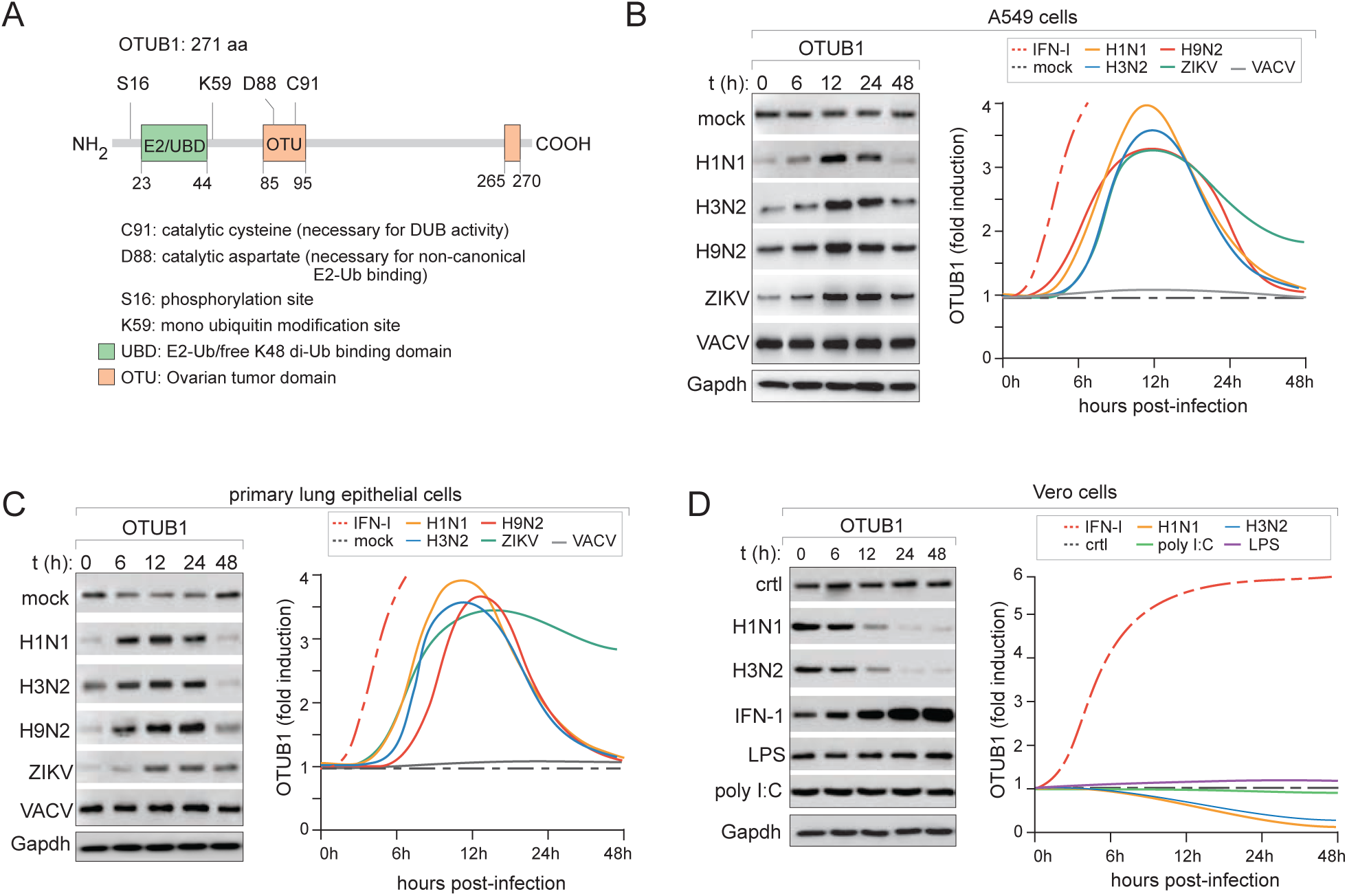
OTUB1 is interferon-inducible and degraded upon influenza infection. **(A)** Schematic of domain architecture of OTUB1 showing its catalytic site, OTU-domain and UBD/E2-binding region, with specific mutations in residues that have been used in the study **(B)** Steady-state levels of OTUB1 in A549 cells infected with different viruses (*left panel*). Lysates collected at indicated time points post infection were immunoblotted with antibodies against endogenous OTUB1. Densitometric analyses of protein expression was performed with ImageJ and normalised to 0 time point. Curve fitting was performed with data from atleast three independent experiments (*right panel*). (**C**) Steady-state levels of OTUB1 in primary lung epithelial cells stimulated with different viruses and assessed as in (B). Curve fitting was performed with data from atleast three independent experiments. **(D)** Steady-state levels of OTUB1 in Vero cells deficient in interferon production stimulated with either Influenza A viruses or ligands alone; LPS to stimulate TLR4 and poly I:C for RIG-I.

OTUB1 expression levels increased as a function of time and peaked ~12 hours post infection in multicycle IAV-infections. This was followed by degradation at later time points, a phenomenon recapitulated in both A549 and primary lung epithelial cells, with all IAV strains (Figure 2B, 2C). To uncouple its induction from degradation, we performed infections in Vero cells deficient in interferon production, where IAV-dependent degradation of OTUB1 was more apparent (Figure 2D). Vero cells treated with exogenous IFN-β displayed a significant increase in the expression of OTUB1, but not its degradation, suggesting that the latter phenomenon was very likely virus-mediated. Stimulation with other ligands (LPS, poly I:C) did not have a significant effect on OTUB1 protein levels in Vero cells deficient in transcriptional induction of IFN-I (Figure 2D).

### OTUB1 is redistributed to mitochondrial membranes and interacts with RIG-I during infection

To establish the subcellular distribution and interactors of OTUB1 in IAV-infected cells, we measured its localisation by confocal imaging, and by proximity-based labelling. In uninfected cells, OTUB1 resided largely in the nucleus; upon infection OTUB1 relocalised from the nucleus to the cytosol and mitochondrial membranes. A significant fraction colocalised with HIF1A, a mitochondrial marker, specifically in infected cells but not in the mock (Figure 3A). Phosphorylation-defective mutants of OTUB1 (Herhaus et al., 2014), S16A (phospho-deficient) and S16E (phosphomimetic), were enriched in the cytosol and nucleus respectively, whereas C91S - the catalytically dead variant, and the D88A mutant - defective in binding E2 enzymes, displayed a similar intracellular distribution as that of the wild-type (**Figure S2**).

**Figure 3.**
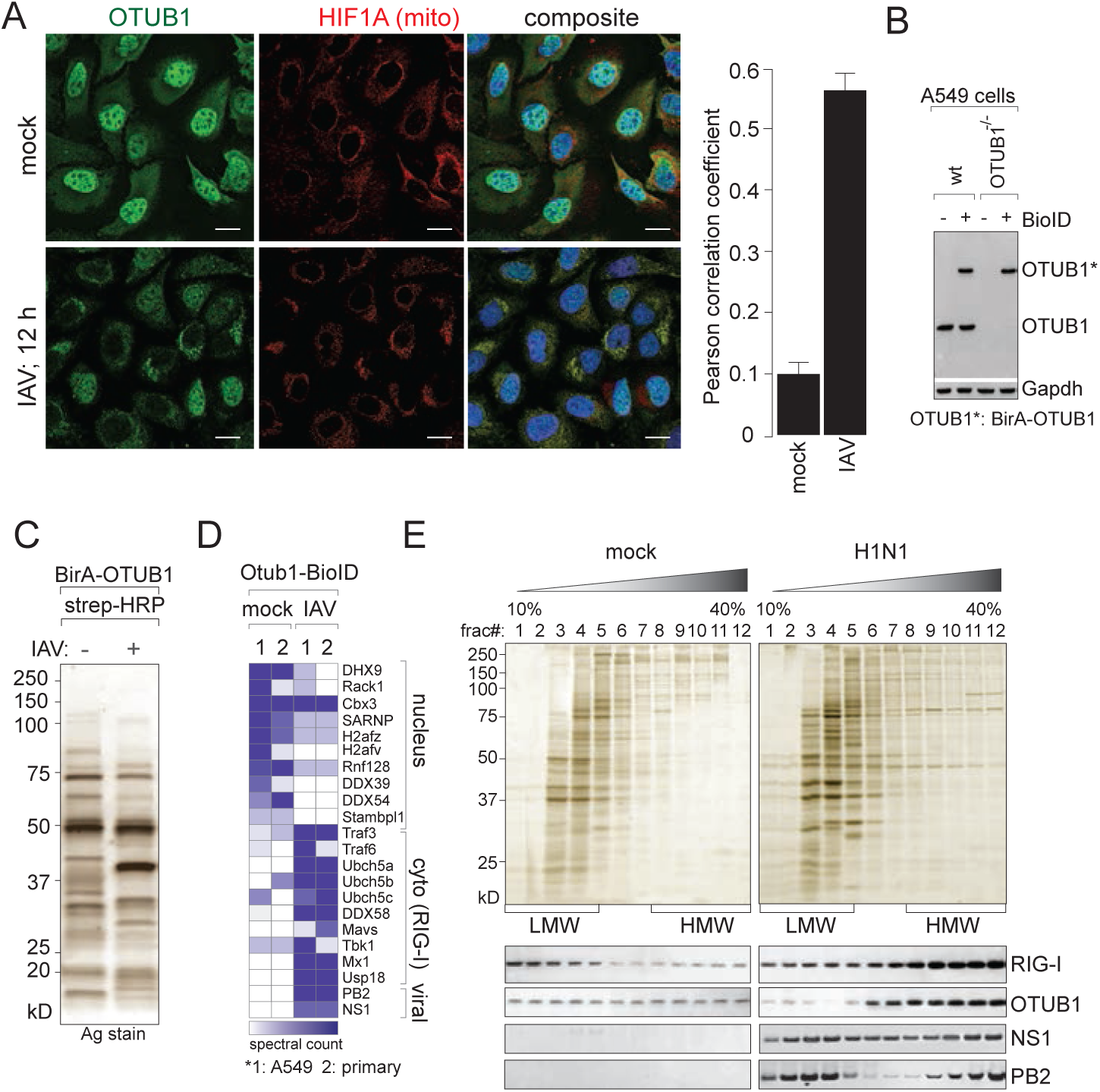
OTUB1 is distributed to the cytosol and mitochondrial membranes upon infection. **(A)** OTUB1 distribution (in green) and mitochondria (in red) visualised in mock-infected and IAV-infected samples using confocal imaging. *Scale bar; 10 µm*. Images representative of >3 independent experiments. Peason’s correlation coefficient was determined to quantitate colocalisation of OTUB1 with that of HIF1 as a mitochondrial marker (*right panel*) **(B)** Stable expression of BioID-OTUB1 construct in either wild-type or OTUB1^-/-^ A549 cells was verified by immunoblotting. **(C)** Silver stained gel of immunoprecipitated material captured on streptavidin beads from mock and IAV-infected A549 cells stably transduced with BioID-OTUB1. 2 mm slices on entire lanes were processed and injected into a Lumos Orbitrap mass spectrometer for peptide identification. **(D)** Spectral counts were determined for candidates identified by mass spectrometry and rank ordered with a threshold of 5-fold difference between mock and infected samples. Significant hits were categorised into GO terms using Ingenuity Pathway Analyses. Two independent biological replicates from different cell types (A549 and primary lung epithelial cells) were performed for MS analyses. **(E)** Lysates prepared from mock and IAV-infected cells were separated by velocity sedimentation on linear glycerol gradients (10%-40%); fractions collected were resolved by SDS-PAGE and detected by silver staining (*upper panel*). Migration characteristics of RIG-I, OTUB1, and IAV proteins NS1 and PB2 from mock and IAV-infected cells were determined by immunoblotting in the collected fractions; HMW: high molecular weight species, LMW: low molecular weight species were calibrated by internal markers.

To identify its interactors in mock and IAV-infected cells, we expressed a BioID-OTUB1 construct in the wt or OTUB1^-/-^ background, detected by immunoblotting (Figure 3B). Mock- or IAV-infected BioID-OTUB1 cells were cultured in biotin-supplemented medium. Biotinylated proteins were captured by streptavidin beads, resolved by SDS-PAGE and identified by LC-MS/MS to measure peptide abundances (Figure 3C, 3D). Identified candidates were sorted by functional enrichment of pathways using Ingenuity Pathway Analyses. The isolated candidates indicated that OTUB1 preferentially interacted with components of the RIG-I signalling pathway in the IAV-infected samples as compared to mock, where it primarily interacted with RNA helicases in the nucleus. Among the viral factors, OTUB1 interacted with PB2 as previously reported (Biquand et al., 2017), and NS1. To confirm MS data, we separated lysates from mock- and IAV-infected cells by velocity sedimentation on glycerol gradients, where OTUB1 co-migrated with high molecular weight RIG-I oligomers in IAV-infected, but not in mock-infected cells (Figure 3E). Interestingly, while IAV NS1 was distributed uniformly through the gradient, PB2 displayed two peaks, with one coinciding with the oligomeric RIG-I, which presumably corresponds to the mitochondrial fraction of PB2. These data suggest that OTUB1 is recruited to mitochondrial membranes where it interacts with active RIG-I tetramers, supporting its role in the RIG-I signaling pathway.

### OTUB1-deletion results in defective NFκB and IRF responses during RNA virus infections

To determine the functional impact of OTUB1 on RIG-I signaling, we generated CRISPR/Cas9-mediated OTUB1^-/-^ A549 cells. Single clones selected after serial dilution were expanded to verify deletion by immunoblotting. Wild-type OTUB1 was then reconstituted back into the deletion background (rOTUB1) (Figure 4A; *left panel*). In parallel, we generated OTUB1^-/-^ A549 cells expressing a dual reporter for IRF and NFκB activities (A549^dual^), measured via luciferase and secreted alkaline phosphatase respectively. Deletion of OTUB1 was created in either the wild-type or a RIG-I^-/-^ background in these reporter cells (Figure 4A; *right panel*).

**Figure 4.**
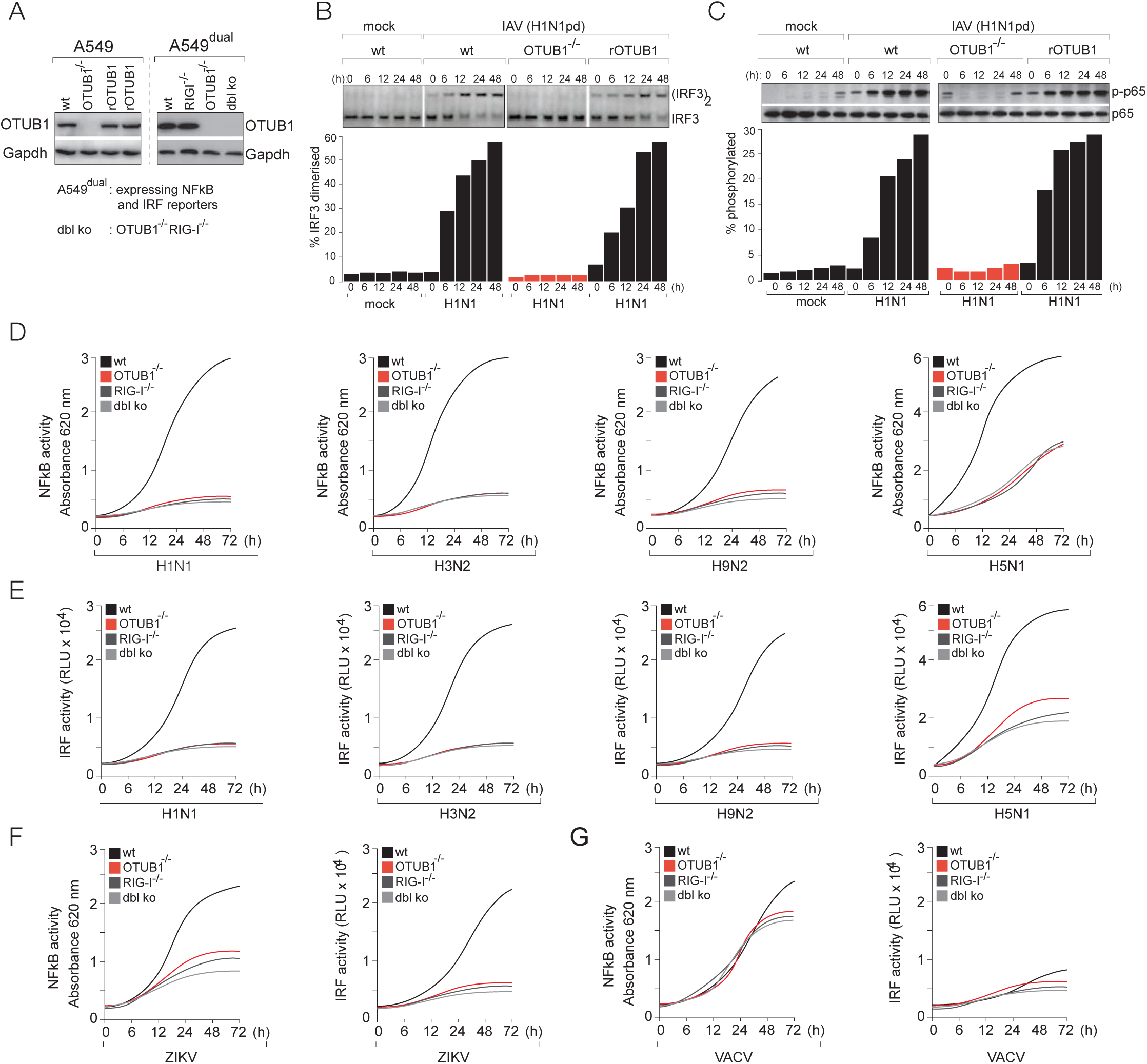
OTUB1-deficiency blocks NFκB and IRF activation. (**A**) Clonal isolates of OTUB1 knock out generated by CRISPR/Cas9 and those reconstituted with wild-type OTUB1 in A549 cells; lysates were used for immunoblotting (anti-OTUB1) to verify deletion and expression (*left panel*). Deletion of OTUB1 in A549 cells expressing a dual reporter for NFκB and IRF activities, either wild-type or harboring a deletion of RIG-I (*right panel*) **(B)** Activation of IRF3 was measured in mock and IAV-infected cells, either wild-type, OTUB^-/-^, or reconstituted with wild-type OTUB1 (rOTUB1). Formation of IRF3 dimers was resolved by native PAGE. **(C)** Activation of NFκB was similarly measured in mock and IAV-infected cells (wild-type, OTUB^-/-^, or rOTUB1) by appearance of phosphorylated p65. Images in (B) and (C) are representative of atleast 3 independent biological replicates. **(D)** Activation of NFκB in cells (wt, OTUB1^-/-^, RIG-I^-/-^, or combined deletion of OTUB1^-/-^ and RIG-I^-/-^), infected with different strains of IAV was measured by secreted alkaline phosphatase measured by absorbance at 620nm **(E)** Activation of IRF3 in cells (wt, OTUB1^-/-^, RIG-I^-/-^, or combined deletion of OTUB1^-/-^ and RIG-I^-/-^), infected with different strains of IAV was measured by luciferase luminescenc **(F)** NFκB and IRF3 activities in ZIKV-infected cells **(G)** NFκB and IRF3 activities in VACV-infected cells, measured as in (D) and (E). Curve fits were generated with data from atleast 3 independent experiments.

Infection by RNA viruses and subsequent RIG-I signaling is accompanied by activation of NFκB and IRF3, which can be detected by phosphorylation of the p65 subunit of NFκB and dimerisation of IRF3. To monitor their activities in OTUB1^-/-^ and rOTUB1 cells, we measured appearance of IRF3 dimers (Figure 4B) and phosphorylated p65 (Figure 4C). In both instances, OTUB1-deficiency resulted in loss of these activities but were rescued in the rOTUB1 cells (Figure 4B, 4C).

To establish how universal this effect was, we infected A549^dual^ cells (wt, OTUB1^-/-^, RIG I^-/-^ or OTUB1^-/-^RIG I^-/-^ double knock-out) with different strains of influenza A virus, including an unrelated RNA virus (Zika) and a DNA virus (Vaccinia) in the sample set. Supernatants collected at different time intervals were quantitated for NFκB activity by measuring absorbance of secreted alkaline phosphatase at 620nm. For all strains of influenza as well as for poly I:C treatment, a significant reduction of NFκB activity was recorded in OTUB1^-/-^ cells - comparable to RIG-I^-/-^ cells. No additive effect was noted in the double knock-out cells, suggesting that they most likely function in the same pathway (Figure 4D, **S3A**).

In parallel, we quantitated IRF activity in the same set of A549^dual^ cells as described above using a luciferase read-out. As with NFκB activation, OTUB1^-/-^ cells displayed significantly diminished IRF activity upon infection with different strains of influenza, as well as with poly I:C (Figure 4E, **S3A**). On the other hand, deficiency of OTUB1 did not have any effect on LPS-depedent activation of NFκB or IRF (**Figure S3B**). Collectively, our results indicate that OTUB1-deficiency inhibits both NFκB and IRF activities, suggesting that it acts upstream of these transcription factors in the RIG-I signalling cascade. OTUB1-deficiency had a significant impact on IRF activity with Zika virus too, supporting the role of RIG-I in viral sensing (Figure 4F); however, no significant effect was detected with Vaccinia infection, since the predominant viral sensing and immune activation to DNA viruses is cGas/STING dependent (Figure 4G).

### The RIG-I signalling complex disassembles in the absence of OTUB1

In uninfected cells RIG-I remains folded with its CARD domains in an auto-repressed inactive comformation; upon infection, viral dsRNA binding exposes the CARD domains which interact with K63 polyubiquitin chains and a multitude of effector molecules for optimal signal transduction in the RIG-I-MAVS-TBK1-IRF3 axis as shown in the schematic (Figure 5A). To determine whether formation of the RIG-I signaling complex requires OTUB1, we transduced myc-RIG-I into RIG-I^-/-^ A549 cells, which either had a deletion of OTUB1 or expressed a wild-type copy, and infected with H1N1(pdm). Most of the signaling molecules displayed enhanced expression in lysates from IAV-infected cells as compared to mock (Figure 5B). While all of them (TRIM25, Riplet, MAVS, TRAF3, TRAF6, TBK1) co-precipitated with myc-RIG-I from cells expressing OTUB1 (either wild-type cells or reconstituted), this failed in the absence of OTUB1 (Figure 5C; *left panel*). In accordance with this effect, K63 polyubiquitin chains could be captured with RIG-I in wild-type cells or those reconstituted with OTUB1, but failed to co-purify in OTUB1^-/-^ cells. We further investigated this complex formation by sedimentation over glycerol gradients. Lysates from IAV-infected wild-type or OTUB1^-/-^ cells were separated by velocity sedimentation over 10-40% linear glycerol gradients. Fractions collected were immunoblotted for RIG-I and associated partners. Most of the signaling components co-migrated with high molecular weight RIG-I in wild-type cells, comprising active tetrameric complexes, but not in OTUB1^-/-^ cells (Figure 5D). This effect extended to MAVS oligomerisation. Although its expression was unaffected in OTUB1^-/-^ cells, multimerisation of MAVS was abolished in OTUB1^-/-^ cells but rescued in those expressing wild-type OTUB1 (Figure 5E). These data indicate that in the absence of OTUB1 RIG-I fails to form active oligomeric structures, thereby impeding recruitment of the signaling complex and IRF3 dimerisation.

**Figure 5.**
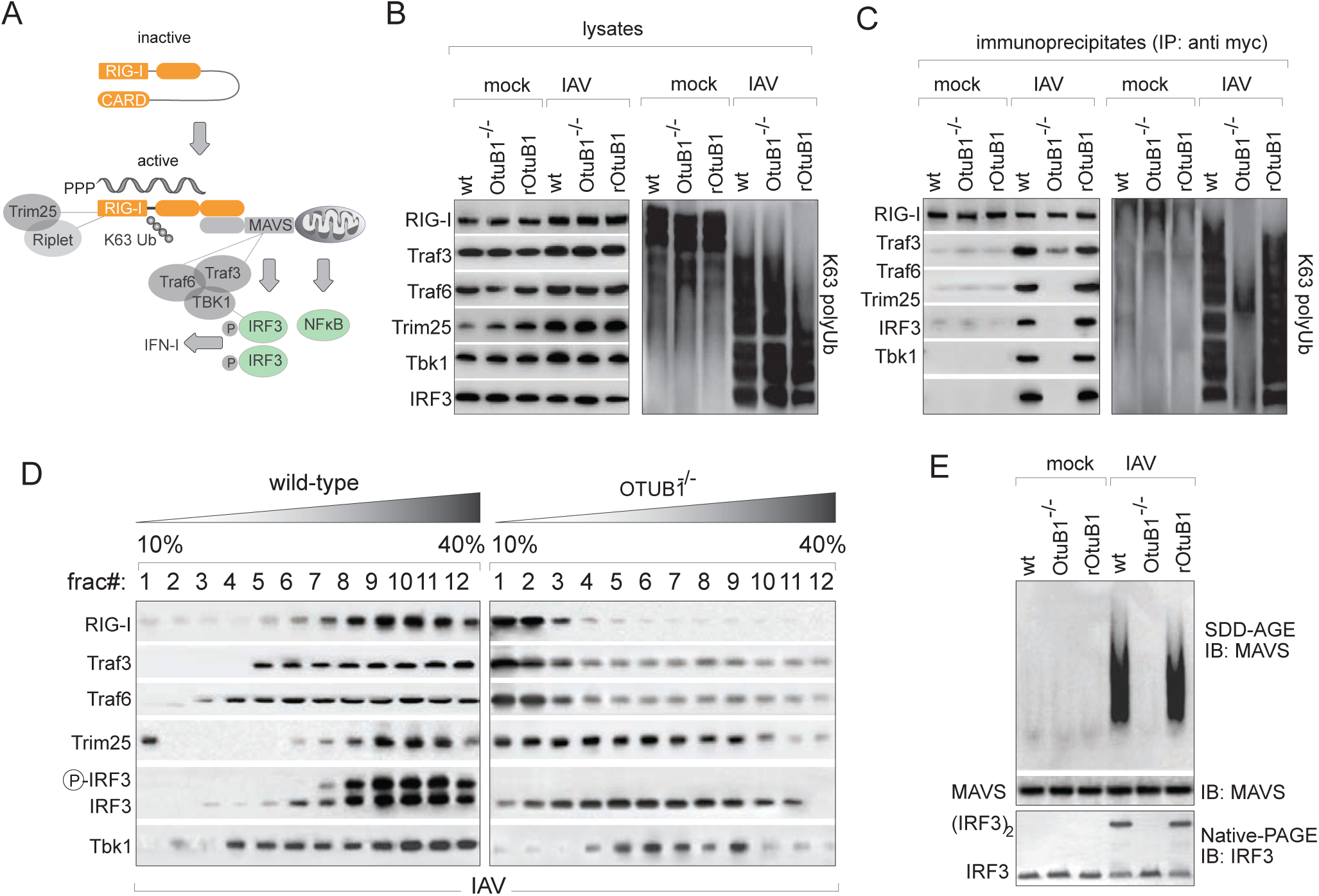
Disassembly of the RIG-I signalling complex in the absence of OTUB1. **(A)** Schematic of the activated RIG-I complex and known interactors of the signaling pathway **(B)** A549 cells stably transduced with myc-RIG-I were genome edited to express either wt, OTUB1^-/-^ or rOTUB1, and exposed to mock or IAV-infection. Signaling molecules were detected by immunoblotting in lysates prepared from mock and IAV-infected cells (*left panel*); presence of K63 polyubiquitin chains in the lysates was measured by anti-K63 antibodies (*right panel*). **(C)** Immunoprecipitation of RIG-I using anti-myc from mock- and IAV-infected cells described in (B) was followed by resolving eluted material by SDS-PAGE and immunoblotting for co-purifiying components (*left panel*) and K63 ubiquitin chains (*right panel*) **(D)** Lysates from wild-type or OTUB1^-/-^ cells infected with IAV was resolved by velocity sedimentation on glycerol gradients (10% - 40% (w/v)); fractions collected were desalted, resolved by SDS-PAGE and visualised by immunoblotting for components of the RIG-I complex. **(E)** Lysates from (B) were also resolved by semidenaturing agarose gels (SDD-AGE) to visualise MAVS oligomerisation in mock and IAV-infected cells (*upper panel*), SDS-PAGE to measure total MAVS levels (*middle panel*), and native-PAGE to measure IRF3 dimerisation (*lower panel*).

### In vitro reconstitution of OTUB1-dependent IRF3 dimerisation activity from mitochondrial and cytosolic fractions

OTUB1 belongs to the OTU-family of cysteine proteases and specifically removes K48 linked polyubiquitin chains. It also functions non-catalytically by binding E2 enzymes that can either inhibit ubiquitylation or stimulate K48 DUB activity, depending on the concentration of free ubiquitin (Wang, 2012). To determine the mechanism of OTUB1-dependent regulation of the RIG-I pathway in IAV-infected cells, we reconstituted IRF3 dimerisation in a cell free system composed of purified RIG-I, mitochondria, [^35^S]IRF3 and cytosol preparations from cells expressing OTUB1 variants (Figure 6A). RIG-I purified from either poly I:C treated or IAV-infected cells, but not from untreated or mock-infected controls, could support IRF3 dimerisation when incubated with cytosol and ATP (Figure 6B). Using this assay, cytosolic preparations from OTUB1^-/-^ cells, or those expressing OTUB1 variants were tested for their ability to dimerise IRF3 (Figure 6C, 6D). Cytosol from OTUB1^-/-^ cells failed to induce IRF3 dimerisation. While cytosol expressing wild-type or S16A (cytosolically enriched) could rescue this defect, those expressing mutants defective in forming the E2 repressive complex (D88A) or restricted to the nuclear fraction (S16E) failed to induce IRF3 dimerisation in the in vitro system. Interestingly, the catalytically dead mutant C91S was able to partially support IRF3 dimerisation, suggesting that either the enzymatic activity of OTUB1 is not required, or that redundancy with other cytosolic DUBs can partially rescue the defect. Collectively these data indicate that the non-canonical function of forming the E2-repressive complex, rather than the enzymatic activity of OTUB1 has a greater impact on IRF3 activation. This phenomenon was recapitulated with RIG-I purified from either poly I:C-treated or IAV-infected cells, but not from untreated or mock-infected cells, indicating that the active conformation of RIG-I is necessary for triggering IRF3 dimerisation (Figure 6C, D).

**Figure 6.**
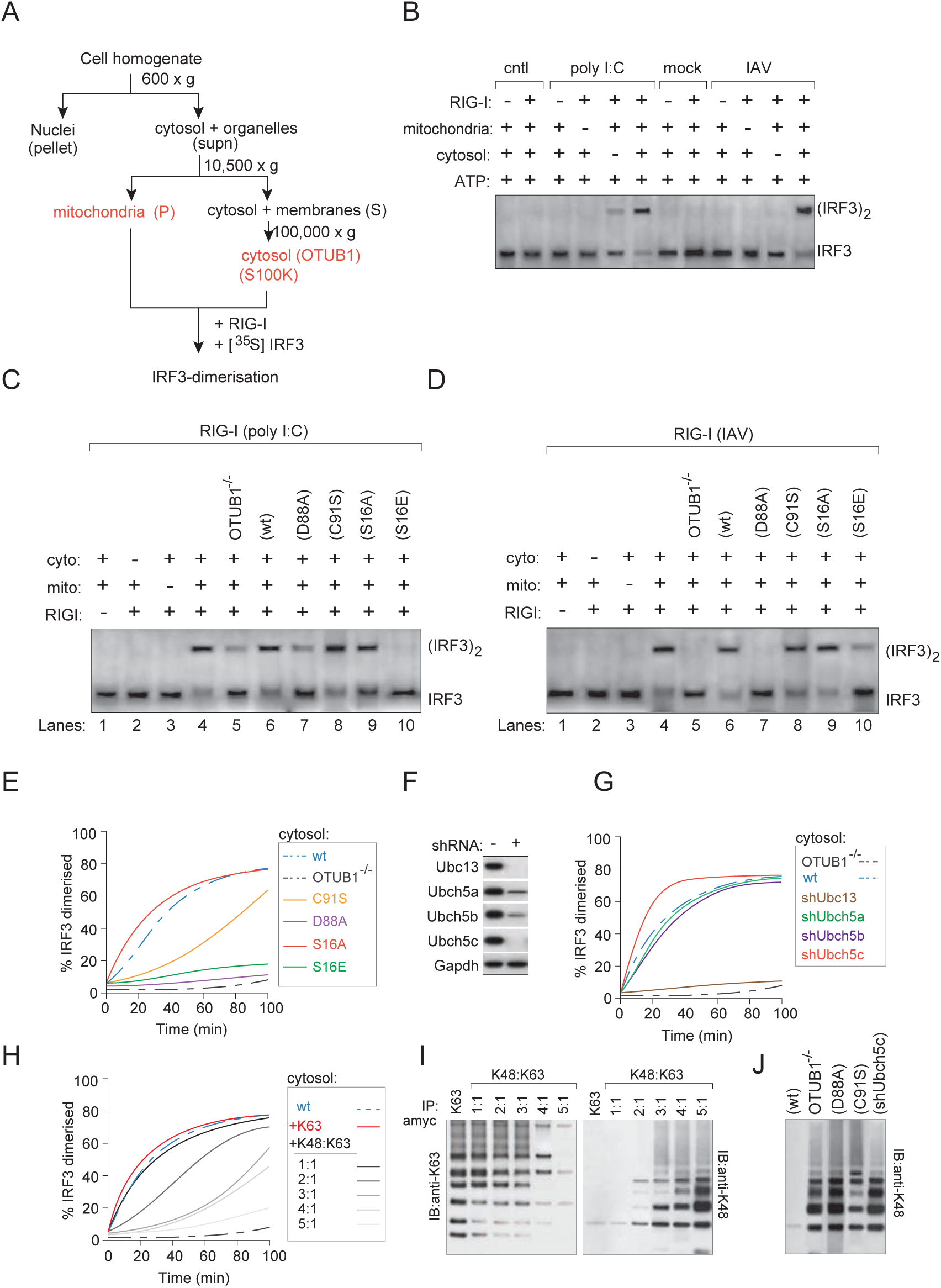
In vitro reconstitution of OTUB1-dependent IRF3 dimerisation. **(A)** Schematic for purification of cytosol and mitochondrial membranes for reconstitution assay. **(B)** IRF3 dimerisation was reconstituted with myc-tagged RIG-I isolated from either IAV-infected or poly I:C treated cells, purified with c-myc peptide, and compared to mock-infected or untreated cells. The reaction mixture contained mitochondria, cytosolic extracts, ATP, [^35^S]IRF3 and RIG-I. Dimerisation of IRF3 was analysed by native gel electrophoresis after incubating for 120 min. **(C)** Cytosolic extracts were prepared from either OTUB1^-/-^ cells or reconstituted with WT OTUB1, D88A, C91S, S16A, S16E mutants to test for their ability to induce IRF3 dimerisation in the assay described in (B), with RIG-I isolated from poly I:C treated cells. **(D)** Same as (C) except RIG-I was isolated from IAV-infected cells. **(E)** Time courses of IRF3 activation was performed with cytosolic extracts as described in (C) and (D) using the assay described in (B). Quantitation of IRF3 dimers was calculated as a percentage of total, from histogram plots generated by ImageJ densitometric analyses. Curve fitting was performed with data obtained from atleast 3 independent experiments. **(F)** The role of E2 enzymes in OTUB1-dependent IRF3 dimerisation. Depletion of UBC13 and UBCH5(a-c) was performed in A549 cells using shRNA targeting the corresponding enzymes. Cell lysates were immunoblotted with antibodies against the endogenous proteins to confirm their depletion. **(G)** Cytosolic extracts generated from either WT or OTUB1^-/-^ cells, or from those depleted in the E2 enzymes from (F) were reconstituted to measure their ability to support IRF3 dimerisation using the assay described in (E). **(H)** Dependence of OTUB1-dependent IRF3 dimerisation on the ratio of K48:K63 ubiquitin chains. The reaction mixture in the IRF3 dimerisation assay was supplemented with either 10µM K63 polyubiquitin (Ub2-Ub7) alone or with increasing amounts of K48 polyubiquitin (Ub2-Ub7) to vary the ratio, with a constant total concentration of 10µM. **(I)** Polyubiquitylation status of RIG-I in the reaction mix was determined at the end point of IRF3 dimerisation time-courses in (H) by immunoprecipitation on anti-myc antibodies and immunoblotting with linkage specific anti-Ub. **(J)** K48 polyubiquitin association with RIG-I was determined as in (I) in reactions with cytosol preparations from WT, OTUB1^-/-^, D88A, C91S and shUbch5c expressing cells.

To further deconstruct OTUB1-mediated activation of RIG-I, we performed time courses on IRF3 dimerisation with cytosol from OTUB1^-/-^ cells or from those expressing its mutant variants. Rapid IRF3 dimerisation occurred with the wild-type and the S16A mutant as noted above, and not with the S16E and D88A mutant constructs, which are nuclear localised and deficient in E2-binding respectively. Interestingly, cytosol expressing the catalytically dead C91S mutant supported significantly slower kinetics but was able to rescue IRF3 dimerisation at later time points, suggesting that other cytosolic DUBs are able to compensate for OTUB1 enzymatic function (Figure 6E). Collectively, our data indicate that the E2~OTUB1 repressive complex formation is a critical step in RIG-I mediated IRF3 activity.

Among the E2 enzymes, UBC13 and UBCH5(a-c) are known to bind OTUB1. We therefore generated a set of cytosolic preparations from cells depleted in the corresponding E2s (Figure 6F), and reconstituted them in the IRF3 dimerisation assay. While UBC13-deficient cytosol was devoid of activity, UBCH5c deficiency displayed even faster IRF3 dimerisation kinetics than that from wild-type cells (Figure 6G). These data recapitulate OTUB1-dependent IRF3 activity and suggest that formation of a UBCH5c~OTUB1 repressive complex activates the RIG-I signaling pathway. UBC13 itself has been described to function in TRIM25-mediated K63 ubiquitylation of RIG-I (Sanchez et al., 2016); it is therefore not surprising that UBC13 depletion inhibits IRF3 dimerisation. On the other hand UBCH5(a-c) have been proposed to function in concert with RNF125 to catalyse K48 polyubiquitylation and proteasomal degradation of RIG-I (Arimoto et al., 2007). Inhibition of UBCH5c by forming an E2-repressive complex with OTUB1 would therefore prevent RIG-I degradation and enable activation of this pathway.

Our data suggest that OTUB1 prevents accumulation and conjugation of K48 polyubiquitin chains to RIG-I. Which means, that altering the local concentrations of K48 and K63 polyubiquitin would in principle regulate association of RIG-I with K63 polyubiquitin and subsequent IRF3 dimerisation. To test this hypothesis, cytosolic fractions from cells expressing wild-type OTUB1 were supplemented with either K63 alone (Ub2-Ub7) or varying ratios of K48:K63 chains to measure IRF3 dimerisation. In line with our hypothesis, increasing concentrations of K48 polyubiquitin resulted in a systematic reduction in kinetics of IRF3 dimerisation (Figure 6H). At K48:K63 ratios >2:1, the rate of IRF3 dimerisation decreased significantly, indicating that concentrations of free polyubiquitin chains can determine the optimal ubiquitylation of RIG-I and regulate the magnitude of the signal transduction cascade.

To detect RIG-I~K63 association in the conditions described above, we immunoprecipitated myc-RIG-I from the reaction mix and immunoblotted with linkage-specific anti-ubiquitin antibodies. The extent of co-precipitating ubiquitin chains reflected those of the K48:K63 ratios and downstream IRF3 activites (Figure 6I). Furthermore, for all cytosol preparations which displayed attenuated IRF3 dimerisation, an increase in K48 polyubiquitin association with RIG-I was noted as compared to cytosol expressing wild-type OTUB1 (Figure 6J). Our data indicate that high local concentrations of K63 polyubiquitin is maintained by OTUB1 activity to ensure optimal RIG-I dependent immune signaling.

### OTUB1 is targeted for proteasomal degradation by IAV NS1

The reduction in steady-state levels of OTUB1 in IAV-infected cells at later timepoints (Figure 2B-D) suggested that it is most likely targeted for degradation. Among the viral interactors identified in the BioID assay, we identified PB2 and NS1 (Figure 3D), both of which are known to antagonise host immune responses (Gack et al., 2009; Graef et al., 2010; Mibayashi et al., 2007). We transfected PB2 and NS1 proteins of IAV to test which of them mediated OTUB1 degradation. Expression of NS1 alone was sufficient to recapitulate reduced steady state levels of OTUB1 (Figure 7A). This was further validated by imaging viral NS1 and OTUB1 in IAV-infected cells, which displayed mutually exclusive expression characteristics (Figure 7B). To investigate the turnover of the newly synthesised pool of OTUB1, we first performed pulse-chase assays in [^35^S]cysteine/methionine labeled cells expressing either the wild-type or C91S mutant of OTUB1. Cells were infected with either wild-type PR8 IAV or two mutants – one lacking NS1 (ΔNS1) or carrying a mutant NS1, lacking its effector domain (mNS1). At indicated time intervals, OTUB1 was immunoprecipitated and detected by autoradiography. Newly synthesised wild-type OTUB1 underwent rapid degradation in the presence of PR8 (WT), but not with defective NS1. Stability of catalytically dead OTUB1 (C91S) on the other hand was not affected under any condition (Figure 7C).

**Figure 7.**
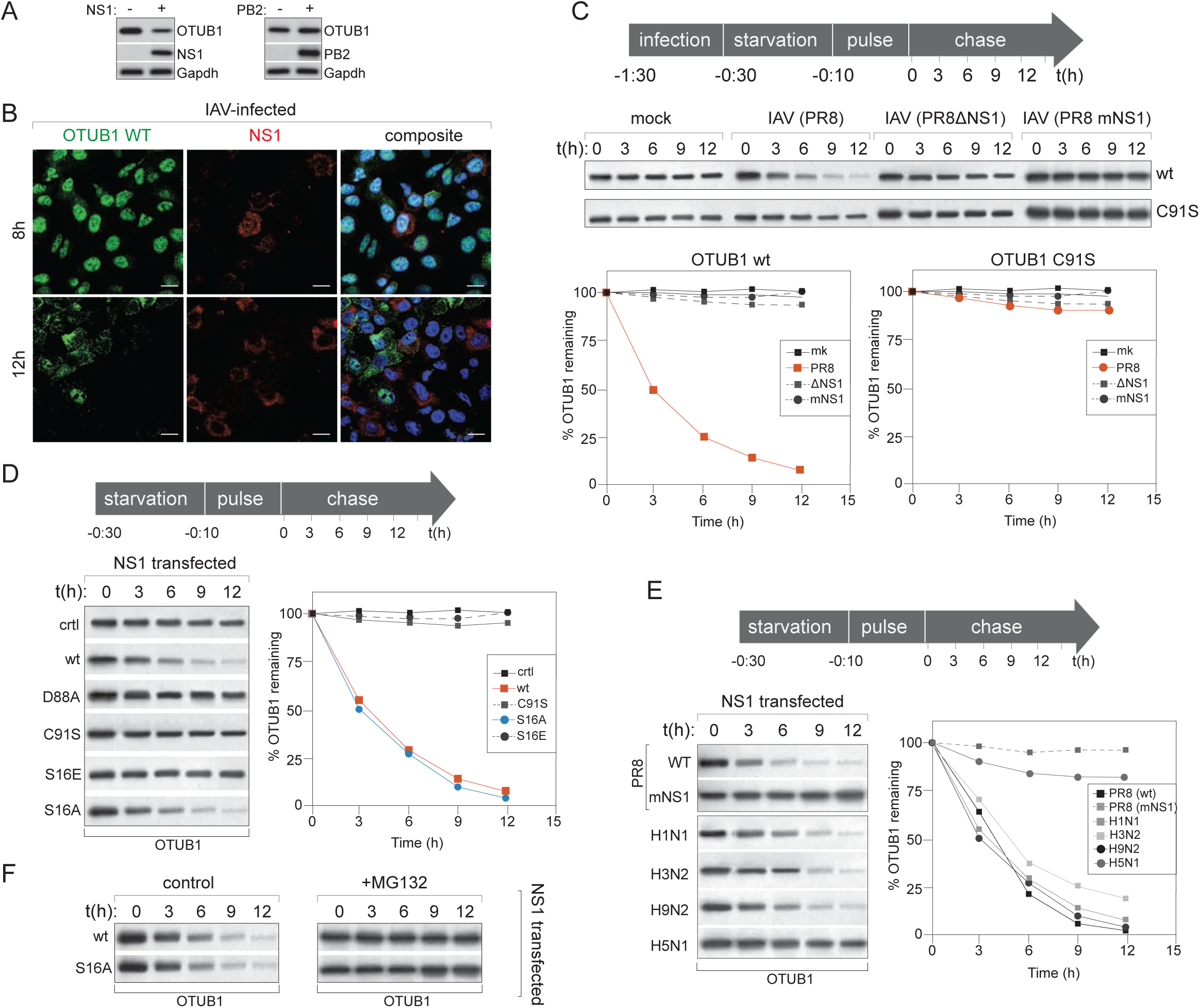
Influenza NS1 triggers proteasomal degradation of OTUB1. **(A)** Steady-state levels of endogenous OTUB1 was measured in A549 cells transfected with either IAV NS1 or PB2. **(B)** Expression of OTUB1 and IAV NS1 are anticorrelated. Confocal imaging was performed on cells infected with IAV (H1N1 pdm). At 8h and 12h post infection, cells were fixed and stained to visualise distribution of OTUB1 and NS1. *Scale bar; 10 µm*. Images representative of >3 independent experiments. **(C)** Wild-type OTUB1 but not C91S catalytic mutant is degraded by IAV NS1. Pulse-chase analyses were performed with OTUB1^-/-^ cells expressing either WT OTUB1 or C91S mutant, and infected with either PR8 (WT), PR8 (ΔNS1) or PR8 (mNS1), which lacks the effector domain. Infected cells were pulsed with [^35^S]cysteine/methionine and chased for indicated time intervals. At each timepoint OTUB1 (wt or C91S) was immunoprecipitated and detected by autoradiography. Amount of OTUB1 remaining was calculated using densitometry as a percentage of total at the 0 time point, normalised to 100%. **(D)** OTUB1^-/-^ cells expressing either the wt or mutant variants were transfected with IAV NS1 and pulse chase assays were performed to measure OTUB1 turnover as described in (B). **(E)** NS1 constructs from different IAV strains as indicated were transfected into cells for pulse chase analyses as described in (B) and (C). **(F)** NS1 from PR8 was transfected into cells expressing either wild-type or S16A variant of OTUB1, and either left untreated or treated with 5µM MG132 for 4 hours to inhibit proteasomal degradation. Pulse-chase analyses of OTUB1 turnover was determined as described in (B) and (C).

To further examine this effect, we transfected NS1 into cells expressing OTUB1 variants, including wild-type, C91S, D88A, S16E and S16A, and compared to control cells transfected with the empty vector control. Both the wild-type and cytosol-localised S19A mutants were rapidly degraded in the presence of NS1, whereas C91S (the catalytic mutant), D88A (deficient in binding to E2) and S16E (nucleus-localised) remained unaffected and stable upon expression of viral NS1 (Figure 7D).

To test whether NS1-mediated degradation of OTUB1 was a universal phenomenon among influenza viruses, we expressed NS1 variants from different strains of IAV. Wild-type NS1 from PR8 was able to rapidly degrade OTUB1, whereas a mutant lacking the effector domain had no effect on OTUB1 turnover. Expression of NS1 from all strains (H1N1 pdm, H3N2, H9N2) except H5N1 were all able to degrade newly synthesised OTUB1 (Figure 7E). Degradation of OTUB1 was blocked upon proteasomal inhibition by MG132 (Figure 7F). Collectively, our data indicate that OTUB1-dependent activation of RIG-I is impeded by the presence of influenza A virus NS1 by facilitating its proteasomal degradation.

## Discussion

RIG-I like receptors (RLR) are critical cytosolic sensors for pathogen-associated molecular patterns (PAMPs), and play key roles in prompting innate immune responses against microbial infection. Viral sensing triggers RIG-I tetramerisation, K63 ubiquitylation and recruitment of signaling adaptor molecules that culminates in NFκB and IRF activation to induce expression of antiviral genes. Not surprisingly, during the course of their co-evolution, viruses have acquired strategies to subvert such immune mechanisms by various means, such as exploiting host factors to their own advantage or triggering their degradation to dysregulate intracellular processes.

For many immune signaling cascades, ubiquitylation is strategically positioned to fine-tune their magnitude and duration, and therefore is also an attractive target for viral subversion mechanisms (Ribet and Cossart, 2010). In this study, we profiled deubiquitylases that are activated upon IAV infection and identified OTUB1 as a critical regulator of RIG-I that is targeted for degradation by IAV NS1. OTUB1 was interferon inducible, associated with the RIG-I signaling complex – specifically the active oligomeric pool at the mitochondrial membranes, and was targeted for proteasomal degradation during IAV infection. Deletion of OTUB1 resulted in impaired IRF3 and NFκB activation - equivalent to that of RIG-I deletion – with all tested strains of IAV, and could be rescued by reconstitution with a wild-type variant into the deletion background. Inhibition of IRF3 resulted from a block in RIG-I oligomerisation, disassembly of the signaling complex, and a subsequent loss of MAVS multimerisation in OTUB1-deficient cells.

To determine the mechanism of OTUB1-dependent RIG-I activation, we adopted a cell free system of IRF3 dimerisation (Zeng et al., 2010), reconstituted from purified RIG-I, mitochondria and cytosol expressing OTUB1 variants. OTUB1 specifically hydrolyses K48 polyubiquitin and has been described by multiple studies to bind and sequester E2 enzymes to inhibit ubiquitylation (Herhaus et al., 2013; Sun et al., 2012). Mutation in its catalytic cysteine (C91S) abolishes its enzymatic function; a D88A mutation blocks its ability to bind to E2 enzymes (Sun et al., 2012), and perturbation in its phosphorylation alters it subcellular distribution; S16A confines it to the cytosol and S16E restricts it to the nucleus (Herhaus et al., 2015). Cytosol fractions from cells deficient in OTUB1 did not support IRF3 dimerisation in the cell-free reconstituted assay. Among cytosol preparations expressing the various mutants, wt OTUB1 was proficient in supporting IRF3 activation, C91S displayed significantly slower kinetics, suggesting that this activity is probably rescued by other cytosolic DUBs; however, with the D88A mutant, IRF3 activity was completely abolished. The cytosolic S16A mutant performed as well as the wild-type, whereas nucleus-localised S16E failed to rescue the defect. Collectively these data indicate that both the DUB activity and E2-repressive function of OTUB1 are employed in the RIG-I signaling pathway, with the former being redundant with other cytosolic DUBs. To determine which E2s might function in this cascade, we generated cells depleted of the known E2-interactors of OTUB1, UBCH5(a-c) and UBC13. Depletion of UBC13, which has been described to function in the TRIM25-dependent K63 ubiquitylation of RIG-I resulted in abrogation of IRF3 activity, whereas that of UBCH5c in particular, allowed for even faster IRF3 dimerisation in the reconstitution assay, reminiscent of OTUB1 rescue assays. Our data suggest that OTUB1 exerts a dual mechanism to enable RIG-I activation – first through its conventional deubiquitylating activity, and the second via inhibition of E2 enzymes, particularly Ubch5c, to stabilise the RIG-I complex and inhibiting proteasomal degradation. The two distinct activities are likely regulated by the local concentrations of K48 and K63 polyubiquitin chains; accumulation of K48 polyubiquitin resulted in inhibition of K63 polyubiquitin association with RIG-I, inhibition of IRF3 dimerisation and increased K48 chain association of RIG-I. A combined effect of hydrolysing K48 chains and inhibiting UBCH5c-mediated K48 ubiquitylation of RIG-I would provide tight regulation of RIG-I activation and amplification of signaling cascade upon infection. Not surprisingly, expression of the multifunctional IAV NS1 alone is sufficient to downregulate this phenomenon through proteasomal degradation of OTUB1. The identity of the E3 ligase that is co-opted by NS1 to degrade OTUB1 is currently under investigation.

This study emphasises the intricate and complex dynamic that exists in the utilization of DUBs in the host-pathogen arms race. Several DUBs have been implicated in different signaling events. OTUB1 itself has been reported to play a role in TNFα and TGFβ signaling cascades. In addition, it also functions in DNA damage repair and histone modification pathways through inhibition of UBC13 and subsequent K63-linked ubiquitylation. A mechanistic understanding of its function in the context of infection provides fundamental insights into regulation of cell signaling and viral evasion strategies.

## Acknowledgements

This work was funded by Research Grants Council (GRF grants 17117914 and 17113915), and partially supported by Health and Medical Research Funds (16150592), theme based research grant from the Research Grants Council (Project No. T11-705/14N), PTR (546) from Institut Pasteur and contribution from BNP Paribas. SS is supported by the Croucher Foundation. The authors acknowledge Michael Weekes and Benedikt Kessler for suggestions with mass spectrometry analyses.

## Author contributions

SS was responsible for the overall design of the study; ASJ, EB, RM, CKM, HHW, AQ, and QWT performed the experiments; AC, RM, LLMP, CD and AG-S were involved in generation and characterisation of constructs, recombinant viruses and cell lines critical to the study; all authors commented on the manuscript.

## STAR methods: Contact for resource and reagent sharing

Further information and requests on reagents and resources should be directed to and will be fulfilled by the lead contact, Sumana Sanyal.

## Experimental models

### Cell Lines

The following cell lines were used in this study: A549 cells (human; sex: male; ATCC-CCL-185), A549^dual^ (human; sex: unspecified; Invivogen a549d-nfis), HEK293T (human; sex: unspecified) and Vero E6 cells (*Cercopithecus aethiops;* sex: unspecified) were obtained from commercial sources noted in the Key Resources Table and gender of the cell line was not a consideration in the study. Cells were maintained in DMEM or EMEM supplemented with 10% FBS as specified by the manufacturer. Primary lung epithelial cells (human; sex: batch-specific; ATCC-PCS-300-010) were maintained according to the manufacturer’s protocol.

### Virus stocks

A/WSN/33 (H1N1), A/Oklahoma/309/06 (H3N2), A/Oklahoma/323/03 (H3N2), A/Hong Kong/1073/99 (H9N2), A/H1N1 09 (H1N1 pandemic), A/Hong Kong/483/1997 (483/H5N1) influenza viruses were propagated in Madin Darby canine kidney (MDCK) cells in minimum essential medium (MEM) supplemented with 0.3% bovine serum albumin (BSA) (Sigma-Aldrich) in the presence of 1 μg/ml tosylphenylalanyl chloromethyl ketone (TPCK)-treated trypsin (Thermo Scientific), 1% (v/v) penicillin/streptomycin (P/S; Sigma-Aldrich), and supernatants from the virus culture were harvested three days post-infection. In order to maintain the homogenicity of the viruses, they were propagated with limited passage number and seed stocks of viruses were prepared for future propagation. Virus stock was aliquoted and stored at −80°C for titration and storage. Viral titer was calculated using plaque assay. ZIKV (MR766) and *Vaccinia* virus strain WR were titrated by determining the tissue culture infective dose 50% (TCID_50_/ml) in Vero E6 cells challenged with 10-fold serial dilutions of infectious supernatants for 90 min at 37°C.

## Method details: Virus infections

A549/ Vero/ 293T cells were seeded in 6-well plates with the seeding density of 0.8 × 10^6^ cells per well one day before the experiment. Cells were infected with viruses at a multiplicity of infection (MOI) of 4 (single cycle) or 0.01 (multi-cycle growth). After an hour of adsorption, the viral inoculum was discarded, and infected cells were then washed with PBS and cultured in Opti-MEM added with TPCK-treated trypsin (0.5-1 µg/ml) in a 37 °C incubator. Supernatants, cell lysates and RNA were harvested at indicated time points, cleared by centrifugation and stored at −80°C until ready to use.

### RT-qPCR assay to measure virus infection

Total RNA was extracted from cells using minibest universal RNA extraction kit (Takara) according to the manufacturer’s manual. 1µg of total RNA was used for the following reverse transcription assays. For the quantification of vRNA, 10 µM of vRNA specific primer complementary to the 3’ end of vRNA and 10 µM of β-actin specific primer complementary to the 3’ end of β-actin gene was used together with SuperScript III Reverse Transcriptase (Invitrogen); while for mRNA measurement 500 ng of oligo-dT primer was used in another reverse transcription reaction. To determine the gene copy number of vRNA and mRNA from M gene, a SYBR Green based real-time PCR method (Roche) was used, and the number of β-actin mRNA was used to normalize the total RNA concentration between different samples, as described previously (Hui*, et al.*, 2009). The PCR experiments were performed using the LightCycler system 480 (Roche). A reaction mix of 20 µl was composed of 1 µl of each gene-specific primer at 10 µM, 10 µl of SYBR Green master mix, 5 µl of 10-fold diluted cDNA and 3 µl of distilled water. The amplification program was as it follows: 95℃ for 5 min, followed by 45 cycles of 95℃ for 10 sec, 60℃ for 10 sec and 72℃ for 10 sec. The specificity of the assay was confirmed by melting-curve analysis at the end of the amplification program. The primers for M gene and β-actin detection are described in the Key Resources table.

### Plaque assays

MDCK cells were seeded in 6-well plates with a seeding density of 1 × 10 ^6^ per well. The following day confluent MDCK plates were washed twice with PBS. 100μl of 10-fold serial dilutions of supernatants were added to monolayers for adsorption. After one h of incubation at 37 °C, the viral inoculum was discarded, and overlaid with MEM containing 1% agarose and 1μg/mL TPCK-treated trypsin, and incubated inverted at 37°C for 3 days. Cells were fixed with 10% formalin in PBS overnight. and stained with 1% crystal violet solution containing 20% ethanol. Plaques were counted and recorded after drying the plates.

### Minigenome viral replicon assay

Cells were transfected in 12-well plates with plasmids encoding PB1, PB2, PA, and NP proteins (320 ng of NP, 160 ng [each] of PB1 and PB2, and 40 ng of PA), together with a plasmid expressing negative-sense firefly luciferase using the Lipofectamine 2000 transfection reagent (Invitrogen), and incubated at 37°C. Twenty hours after transfection, cells were lysed with 300 μl of passive lysis buffer (Promega), and firefly luciferase activity was measured using a luminometer.

## Screen for isolation of deubiquitylases

~1 x 10^7^ cells were detached from 10 cm dishes by brief trypsinization, washed once with Hank’s balanced salt solution (HBSS) and resuspended in 100 µl (HBSS) on ice. PFO was added to cells to a final concentration of 100 nM and maintained on ice for 5 min. The reaction mix was supplemented with an ATP regenerating mix, 10µM HA-Ubvme or TAMRA-Ubvme and protease inhibitor cocktail, and transferred to 37°C for 20 min. The reaction was terminated with lysis buffer. TAMRA-Ubvme reactive DUBs were resolved by SDS-PAGE and visualised by fluorescence scanning, whereas HA-Ubvme labeled DUBs were first isolated on anti-HA beads, resolved by SDS-PAGE and visualised by silver staining.

## Immunoprecipitation and mass spectrometry

Large-scale immunoprecipitations were performed on conjugated HA-beads (for ubiquitylated substrates) and streptavidin beads (for BioID). Entire lanes were sliced into 2 mm sections, and were further processed in 50% water/methanol. Samples were trypsinized and subjected to an Lumos Orbitrap mass spectrometer for identification of candidates. MS/MS spectra were analysed using Sequest algorithm. The target sequences comprised the human protein repository of the Uniprot database and protein sequences corresponding to the Influenza A PR8, WSN and H1N1 pandemic strains. Decoy sequences were obtained upon reversing the orientation of target sequences. Allowed criteria for searches required trypsin cleavage (two missed cleavages allowed), peptide mass tolerance of 20p.p.m., variable oxidation of methionine residues, and static carbamylation modification of cysteine residues. Peptide-spectrum matches were determined with estimated false discovery rate of 1%. Spectral counts for each condition were combined at a protein level and normalized by protein length to infer protein abundances and intensities in each case. Identified hits were further categorized into different biological pathways using Ingenuity Pathway Analyses software.

## Isolation of OTUB1 interactors in mock and virus infected cells

### BioID assay

~10^6^ A549 cells expressing BirA-OTUB1 were cultured in media supplemented with 1µg/ml doxycycline and 50µM biotin. Cells were lysed in 1 ml lysis buffer (50 mM Tris, pH 7.5, 150 mM NaCl, 5mM EDTA, 1 mM DTT, 0.5% TX-100 and 1X complete protease inhibitor cocktail [Roche]). Lysates were incubated with 500 µl of Neutravidin beads with end-over-end rotation for 1 hour at 4°C. Bound material was saturated with biotin. 10% of the sample was reserved for Western blot analyses. Samples reserved for analyses by mass spectroscopy were washed with 50 mM NH_4_HCO_3_ before subjecting to protease digestion.

## Quanti-blue SEAP Phosphatase assay

QUANTI-blue™ Gold is a two-component kit which contains: - QB reagent and – QB buffer, provided by Invivogen (HK). A standard protocol according to manufacturer’s instructions was followed. The following protocol refers to the use of 96-well plates. 180 μl of QUANTI-Blue™ Solution was dispensed per well into a flat-bottom 96-well plate. 20 μl of sample (supernatant of SEAP-expressing cells) or negative control (cell culture medium) were added and incubated at 37°C for 15 min to 6 h. Optical density (OD) at 620nm was measured using a microplate reader.

## Quanti-luc luciferase assay

QUANTI-Luc™ Gold is a two-component reporter kit which contains: - QUANTI-Luc™ Plus and - QLC Stabilizer, provided by Invivogen (HK). A standard protocol according to manufacturer’s instructions was followed. Quanta-Luc pouches were dissolved in sterile water together with QLC stabilizer. 20 µl of cell supernatant was added to a white opaque 96-well plate. 50 µl of QUANTI-Luc™ Gold assay solution was added to each well. The measurement was carried out immediately using a luminometer.

## Transient transfections and lentiviral transductions

Cells were transfected using either Fugene6 (Roche Diagnostics), TransIT or lipofectamine according to the manufacturer’s instructions. For generation of lentiviral particles, 293T cells were seeded at a density of 4 x 10^6^ cells/dish (10 cm dishes) 24h before transfection (in 12 mL of DMEM 10% FBS, 1% P/S, 1% Hepes). Transfection was carried out with TransIT reagent, following the manufacturer’s instructions. Plasmids VSV-G, PLP1 (GAG/POL), PLP2 and plenti vector (plasmid of transfection, the supernatant was removed with a syringe, filtered on 0.22µm and stored overnight at −80°C. Virus particles collected from supernatants were concentrated by ultracentrifugation (28000 rpm, 4°C). Pellets were resuspended in 100 µl of PBS and left at 4°C for 1h to resuspend the virus. A549 cells were seeded in 10 cm dish with a density of 4 x 10^6^ cells/dish before lentiviral transduction. Transduction was performed using 30 µl of virus/dish (with dropwise distribution). 2 days after, the medium was changed and supplemented with 5 µg/ml of blasticidin. Medium was replaced every three days and selection pressure was kept for two weeks, followed by single clonal expansions, which were verified with western blots for expression of desired protein.

## Pulse-chase analysis of OTUB1 turnover

Pulse chase experiments were performed as previously described (Sanyal et al., 2013) (Zhang et al., 2018). Briefly, ~1 x 10^7^ cells, either mock or IAV-infected were detached by trypsinization and starved for 30 min in methionine/cysteine free DMEM at 37°C prior to pulse labeling. Cells were labeled for 10 min at 37°C with 10mCi/ml [^35^S]methionine/cysteine (expressed protein mix; PerkinElmer) and chased for indicated time intervals. At appropriate time points, aliquots were withdrawn and the reaction was stopped with cold PBS. Cell pellets were lysed in Tris buffer containing TX-100 and pre-cleared with agarose beads for 1 hour at 4°C. Immunoprecipitations were performed for 3h at 4°C with gentle agitation. Samples were eluted by boiling in reducing sample buffer, subjected to SDS-PAGE and visualised by autoradiography.

## Generation of knock-out and knock-down cells

### CRISPR/Cas9 mediated deletion of OTUB1

Potential target sequence for CRISPR interference were found using the rules outlined elsewhere (Mali et al., 2013). Transfection of recombinant plasmid into were subjected to puromycin selection at a concentration of 3 µg/ml for A549 and 4 µg /ml for 293T. After 3 days, medium was replaced with that without puromycin and selected cells were allowed to grow for 1-2 weeks depending on the number of surviving cells. Deletion efficiency was checked in the mixed populations. For clonal expansion, selected cells were seeded in a 10 cm dish at a low density of 20 cells/dish. Pyrex® 8mm cloning cylinders were used to demarcate single colonies and silicone grease was used to create an isolated well. Trypsinised colonies from isolated cylinders were grown in 12 well-plates and verified at passages 1, 3 and 10 for complete deletion of OTUB1.

## In vitro reconstitution of IRF3 dimerisation

Biochemical assays for IRF3 activation with cytosolic extracts and mitochondrial preparations were as described previously (Zeng et al., 2010). Briefly, hypotonic buffer [10 mM Tris-HCl (pH 7.5), 10 mM KCl, 1.5 mM MgCl_2_, 0.5 mM EGTA, and protease inhibitor cocktail] was used to prepare cytosol fractions, and isotonic buffer (hypotonic buffer plus 0.25 M D-Mannitol) was used to prepare mitochondrial membranes. Cells were homogenized in appropriate buffers, and centrifuged at 600 g for 5 minutes to pellet nuclei. Post-nuclear supernatants were further centrifuged at 10,500 x g for 10 minutes to collect mitochondrial membranes in the pellet fraction. Supernatants were subjected to centrifugation at 100,000 g, to separate cytosol fractions from microsomes. Pelleted mitochondrial membranes were washed once with isotonic buffer. For IRF3 pathway activation by RIG-I, the reaction mixture contained 1 mg/ml mitochondrial prep, 3 mg/ml cytosol fractions (from OTUB1-deficient cells expressing OTUB1 variants), 1xMgATP buffer [20 mM Hepes-KOH (pH7.4), 2 mM ATP, 5 mM MgCl_2_], and [^35^S]IRF3. After incubation at 30°C for 2 hours, samples were centrifuged at 20,000 g for 5 minutes and supernatants were subjected to native PAGE to visualise IRF3 dimerisation by autoradiography using

## Glycerol gradient analyses

500 μl lysates from mock or infected cells was loaded onto a 3.8 ml glycerol gradient (10–40% (w/v)) prepared in 1% (w/v) NP-40, 100 mM NaCl, 20 mM Hepes-NaOH pH 7.4. Sedimentation standards (ovalbumin (3.6S), BSA (4.2S), β-amylase (8.9S) and catalase (11.4S)) were analyzed in a parallel gradient. The gradients were centrifuged in a Beckman MLS 50 rotor at 35,000 rpm (~165,000 g_av_) for 20 h at 4°C and fractions (300 μl) were collected from the top. The fractions collected were desalted on Bio-Gel P6 spin columns to remove glycerol, resolved by SDS-PAGE and immunoblotted to visualise separation of RIG-I and OTUB1.

## Immunofluorescence assays

For fluorescence microscopy, A549 cells were seeded the day before the experiment, on slides pre-coated with Poly-L- Lysine (sigma). Seeding density used was 0.3 × 10^6^ per well (in 12 well plates). Cells underwent infection with IAV or IFN type I treatment. Time points were collected by washing the slides twice with PBS, followed by fixation with 4% formaldehyde for 20 minutes and incubation in PBS at 4°C. PBS with 0.05% Triton as permeabilization agent was used for 5 minutes followed by two washes with PBS alone. The slides were blocked with either PBS + 4% FBS or PBS + 4 % NGS, for 30min −1 hour at room temperature. Primary antibodies at their appropriate dilution in PBS was added and incubated for 2 hours at room temperature followed by washing twice with PBS. Secondary antibodies diluted in PBS was dispensed and incubated for 1 hour at room temperature. After two washes with PBS, the slides were stained with freshly diluted DAPI staining (sigma) for 2 minutes and rinsed twice with PBS. Slides were dried and placed on glass slides overlaid with mounting buffer and kept overnight at 4°C before visualising with confocal microscopy.

## Quantification and statistical analyses

For RT-qPCR, fold changes in mRNA expression were determined using the ΔΔCt method relative to the values in control samples as indicated in figure legends, after normalization to housekeeping genes. Results obtained from RT-qPCR and plaque assays are presented as means ± standard deviation (SD), unless stated otherwise. Comparisons between cells were performed using ANOVA, unless stated otherwise. Statistical analyses were performed in GraphPad PRISM 7.0.

