## Supplemental information for "OTUB1 is a key regulator of RIG-I dependent immune signalling and is targeted for proteasomal degradation by influenza A NS1"

| Accession no | Protein name | Mol mass (kDa) | Unique peptides Mock | H1N1 | Mowse score | No of MS/MS queries | Seq coverage |
|---|---|---|---|---|---|---|---|
| **Q96FW1** | Ubiquitin thioesterase protein OTUB1 | 37 | 5 | 21 | 901 | 30 | 52 |
| **Q9UHP3** | Ubiquitin carboxyl-terminal hydrolase 25 | 122 | 9 | 14 | 511 | 22 | 37 |
| **Q96RU2** | Ubiquitin carboxyl-terminal hydrolase 28 | 122 | 6 | 14 | 285 | 5 | 28 |
| **Q9Y614** | Ubiquitin carboxyl-terminal hydrolase 3 | 59 | 12 | 22 | 140 | 8 | 15 |
| **Q92560** | Ubiquitin carboxyl-terminal hydrolase BAP1 | 80 | 10 | 16 | 163 | 20 | 9 |
| **Q13107** | Ubiquitin carboxyl-terminal hydrolase 4 | 108 | 28 | 8 | 57 | 4 | 12 |
| **Q9Y5K5** | Ubiquitin carboxyl-terminal hydrolase isozyme L5 | 38 | 13 | 31 | 177 | 5 | 10 |
| **P15374** | Ubiquitin carboxyl-terminal hydrolase isozyme L3 | 26 | 11 | 31 | 213 | 5 | 8 |
| **Q96G74** | OTU domain containing protein 5 | 61 | 24 | 7 | 154 | 16 | 11 |
| **Q7L8S5** | OTU domain containing protein 6A | 33 | 11 | 20 | 284 | 11 | 7 |
| **Q9NQC7** | Ubiquitin carboxyl-terminal hydrolase CYLD | 107 | 12 | 34 | 41 | 4 | 5 |
| **P21580** | Tumor necrosis factor alpha-induced protein 3 | 90 | 6 | 18 | 58 | 2 | 6 |
| **Q96DC9** | Ubiquitin thioesterase OTUB2 | 27 | 7 | 11 | 44 | 1 | 8 |
| **Q9Y4E8** | Ubiquitin carboxyl terminal hydrolase 15 | 112 | 10 | 19 | 31 | 3 | 10 |

Table 1: Identification of deubiquitylases by HA-Ubvme in primary lung epithelial cells, either mock or H1N1 (pdm) infected.
(Related to Figure 1)

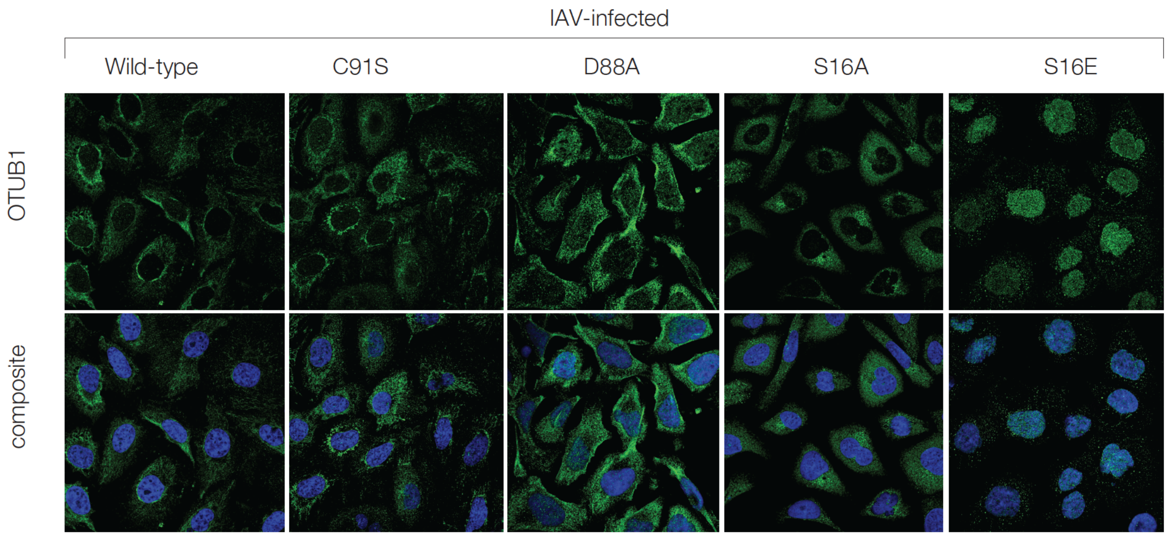


**Figure S2: Subcellular distribution of OTUB1 mutant variants in IAV-infected cells (Related to Figure 3)**

OTUB1$^{-/-}$ A549 cells expressing the different variants of OTUB1 were generated by lentiviral transductions. C91S; catalytic mutant, D88A; deficient in binding to and forming the E2-repressive complex, S16A; phosphorylation deficient and expected to be enriched in the cytosol, S16E; phosphomimetic and expected to be enriched in the nucleus. Subcellular distribution of the panel of OTUB1 mutants were visualised by confocal imaging in IAV (H1N1 pdm) infected A549 cells expressing the OTUB1 variants. Wild-type, C91S and S16A displayed a similar distribution – predominantly present in the cytosolic and mitochondrial compartments. D88A displayed a lack of distribution in the mitochondrial membranes whereas S16E remained confined to the nucleus upon IAV infection.

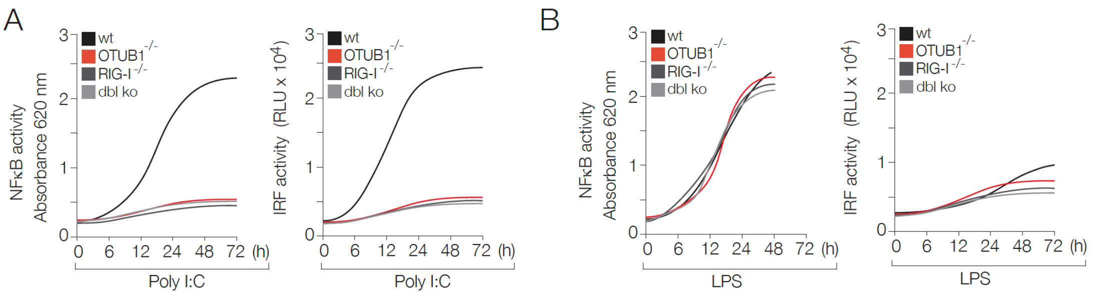


**Figure S3: NFκB and IRF3 activities in Poly I:C and LPS treated cells (Related to Figure 4)**

OTUB1 deletion was generated through CRISPR/Cas9 in A549 cells expressing reporters for NFκB and IRF activities measured via secreted alkaline phosphatase and luciferase respectively. OTUB1$^{-/-}$ cells were generated either in a wild-type background or those harboring a deletion of RIG-I to create a double knockout of OTUB1 and RIG-I. **(A)** The wild-type, OTUB1$^{-/-}$, RIG-I$^{-/-}$, and the double knock-out cells were treated with Poly I:C to stimulate the RIG-I pathway. **(B)** LPS-treated samples were monitored as control. OTUB1$^{-/-}$ cells displayed attenuated NFκB and IRF activities, equivalent to that of RIG-I$^{-/-}$ and the double knock-out cells, when compared to the wild-type cells, when treated with poly I:C. On the other hand, activation of NFκB in LPS treated cells remained insensitive to deletions in OTUB1 or RIG-I or the combined deletion of the two. Since LPS does not trigger IRF activity, it remained at basal levels in all cells, upon LPS treatment.
